# Adolescent peer victimization and subsequent cognitive-affective biases in threat attention, reactivity, and interpretation

**DOI:** 10.1101/2025.07.12.664449

**Authors:** Jens Heumann, Manuel Eisner, Denis Ribeaud, Michael J. Shanahan

## Abstract

**Background:** Peer victimization (PV) in adolescence—verbal, social, and physical aggression—may disrupt normative developmental processes and is associated with psychosocial difficulties. Although previous research has established associations between early adversity and altered attention and cognitive-affective processing, the specific mechanisms of information-processing through which adolescent PV may affect later social functioning remain unclear.

**Methods:** We conducted a counterfactual multimethod study in a sample of young adults (age 22; *n* = 191; 47.1% female), comparing individuals with and without a history of PV between ages 11–20. Specifically we employed: (1) an eye tracking task using a free-viewing paradigm to assess spontaneous threat-related attention; (2) a longitudinal analysis of changes in hostile attribution and reactive aggression using baseline data collected prior to PV exposure and follow-up data in young adulthood; and (3) a morph-based task assessing facial emotion discrimination capabilities between happy and angry expressions.

**Results:** Relative to controls, victims exhibited a vigilance–avoidance attentional pattern and blunted affective responses; they showed reduced initial gaze allocation, repeated returns to angry faces, and a tendency to rate more angry faces as neutral. Over adolescence, a flatter decline in hostile attribution and a steeper drop in reactive aggression led to elevated hostile attribution and reduced reactive aggression.

**Conclusions:** These results suggest a victim profile characterized by impaired attentional processing with biased social threat perception, a more hostile view of the social world, and blunted affect. Our study offers preliminary evidence that adolescent PV may induce enduring changes in social information processing, with implications for maladaptive social cognition and increased risk for psychopathology.

## Introduction

One in three youths worldwide is targeted by peer victimization (PV) (Biswas et al., 2020), an adversity involving verbal abuse, sabotage, exclusion, and violence, which can interfere with their fundamental need for social safety and optimal development (Crick & Dodge, 1994; Slavich, 2020). Persistent psychosocial obstacles—including peer rejection, friendship difficulties, and romantic barriers—are common consequences for adolescents victimized by peers and can ultimately lead to loneliness and deprivation (Coelho et al., 2022; D’Urso et al., 2023; Kochel et al., 2012; K. H. Rubin et al., 2009), with associated effects on stress physiology due to the resulting loss of social status and absence of prestige (Sapolsky, 2004; Slavich, 2020). Unsurprisingly, social anxiety and social withdrawal are about three times more common among victims later in young adulthood (Siegel et al., 2009; Silberg et al., 2016; Stapinski et al., 2014). While childhood adversity is known to affect later social adjustment by altering key domains of biopsychosocial development (Crick & Dodge, 1994; Norman et al., 2012; Samson et al., 2024)—such as attention to threat (Briggs-Gowan et al., 2015; O’Mahen et al., 2015), attribution of intent (Perren et al., 2013), reactive aggression (Buss & Shackelford, 1997; Volk et al., 2022), and facial emotion processing (Jaffee, 2017)—the mechanisms of social information-processing through which adolescent PV may influence these domains have been scarcely examined to date (Kellij et al., 2022).

Attention to threat is often conceptualized as an evolutionary adaptation (Ceccarini et al., 2024; Liu et al., 2021; Schimmack, 2005): besides promoting survival by enabling rapid threat detection, it also heightens sensitivity to status-relevant cues (Gilbert, 2001; Schimmack, 2005). Indeed, healthy prepubertal children focus their attention toward threat, while the ability to suppress excessive threat-related attention typically emerges in adolescence as part of normative cognitive maturation (Ceccarini et al., 2024; Dodge, 2006; Kindt & Van Den Hout, 2001; Rapee et al., 2023). However, adolescents and young adults with elevated anxiety symptoms or trauma—as observed in PV (Pontillo et al., 2019)—may retain exaggerated residual attention to threat (Bishop, 2007; MacLeod et al., 1986; Rapee et al., 2023). Social adversity during this crucial learning phase—such as PV—can disrupt the development of healthy attribution of intent, which may later manifest in ambiguous situations as hostile attribution bias (Craske et al., 2009; Dodge & Coie, 1987). Together, attribution of intent and threat-related attention are thought to be interconnected cognitive-affective domains of social threat processing (Van Bockstaele et al., 2014).

Both attention to threat and hostile attribution bias have been closely linked to reactive aggression (Card & Little, 2006; Crick & Dodge, 1994; Dodge & Coie, 1987; Verhoef et al., 2019; Wilkowski & Robinson, 2008), and hostile attribution bias is a key driver of reactive aggression (Crick & Dodge, 1994). Also, PV has been associated with reactive aggression (Heilbron & Prinstein, 2008); victims may exert forms of retaliation (Camodeca & Goossens, 2005; Herd & Kim-Spoon, 2021) or engage in displacement aggression to compensate for loss of social status (Card & Little, 2006; Sapolsky, 2004).

Lastly, processing of facial expressions—particularly of angry emotions—also continues to develop into adolescence (Ceccarini et al., 2024; Durand et al., 2007; Pfaltz et al., 2019; Pollak & Kistler, 2002). Most of the evidence stems from studies on childhood maltreatment, who report heightened sensitivity to anger cues, which is thought to reflect an adaptive threat-avoidance mechanism (Gibb et al., 2009; Pollak & Kistler, 2002). However, there are other studies reporting reduced sensitivity to negative stimuli, particularly following acute stress (F. S. Chen et al., 2014) or trauma exposure in adulthood (Wormwood et al., 2016). Yet socially anxious individuals may still interpret ambiguous expressions as angry—not due to heightened sensitivity, but to avoid the cost of missing dominant threat and to adopt submissive behavior for safety (Maoz et al., 2016). Whether such social stress—especially from PV—leads to biased interpretation of facial expressions with its psychosocial consequences, however, remains unclear.

Threat-related attention is typically assessed using the dot-probe paradigm, during which emotional and neutral stimuli are briefly presented side by side and then replaced by a probe in one location, which participants localize as quickly as possible while their reaction times are recorded (Lisk et al., 2020; MacLeod et al., 1986; Roy et al., 2008). Reaction times methods, however, have yielded mixed results; some studies report attention toward threat (e.g., MacLeod et al., 1986), while others report attention away from it (Iffland et al., 2019; Lisk et al., 2020). One dot-probe study on children exposed to family violence used eye tracking and found faster first fixations but shorter attentional maintenance on emotional faces, suggesting a vigilance–avoidance pattern (Hoepfel et al., 2022). Unlike reaction-time measures, eye tracking offers a direct, time-resolved measure of visual attention across the full trial with fewer confounds (Armstrong & Olatunji, 2012; N. T. M. Chen & Clarke, 2017; Günther et al., 2021), especially in paradigms with longer viewing times than dot-probe, such as free-viewing tasks (Günther et al., 2021).

Hostile attribution bias and reactive aggression are often measured concurrently using questionnaires in which participants read about provocative but ambiguous social situations and indicate the extent to which they attribute hostile intent and how aggressively they would be likely to respond (Camodeca & Goossens, 2005; Ziv et al., 2013). While most existing work is cross-sectional (Klein Tuente et al., 2019), longitudinal designs may be better suited to gain insights into whether and how adversity contributes to shifts in these patterns over time.

Facial emotion perception is commonly examined using paradigms in which participants rate representations of faces displaying emotional expressions—often artificially blended from two emotions (Ito et al., 2017). Such static images, however, are limited in capturing perceptual biases, as they typically assess sensitivity to a single emotion and do not allow direct comparison between two emotional signals (Claudino et al., 2019; Fu & Pérez-Edgar, 2019; Ito et al., 2017; Zupan & Eskritt, 2024). To remedy this, Ito et al. (2017) had participants morph facial expressions between emotions in real time.

Using a counterfactual approach, this study: (1) assesses threat-related attention bias through eye tracking during a free-viewing paradigm—conceptually akin to the dot-probe task but allowing extended exposure; (2) leverages a rare longitudinal design to evaluate changes in reactive aggression, drawing on pre- and post-PV assessments within individuals to capture intra-individual change; and (3) examines facial emotion discrimination bias using an interactive morphing task, during which participants adjust facial expressions to their perceived neutral point.

## Methods

### Participants

Participants were drawn from the Zurich Brain and Immune Gene Study (z-GIG), a subsample (*n* = 200) of the longitudinal z-proso panel (*n* ≈ 1,500; Ribeaud et al., 2022; z-proso Project Team, 2024), enriched for victimization. z-proso began in 2004 with Zurich primary school children. In 2019, following the eighth wave, the z-GIG study was conducted, incorporating physiological, biological, neuroimaging, and psychological outcomes. The present study focuses on cognitive-affective experiments and the Ambiguous Intentions Hostility Questionnaire (AIHQ; Combs et al., 2007).

The analytic sample comprised *n* = 191 participants (mean age = 21.80; SD = 0.497; female: 47.1%). PV was assessed from the fourth wave (mean age = 10.90; SD = 0.40) to the eighth wave (mean age = 20.31; SD = 0.345) from retrospective self-reports across experienced and exerted social peer adversity across the domains verbal abuse, sabotage, exclusion, violence, among others, resulting in prospectively structured data (Murray et al., 2021). A binary PV variable (23 victims; 12.0%) was derived, as detailed in the Supporting Information. Morphing and questionnaire data were available for all 191 participants, whereas eye-tracking data were available for 179 participants, as data from nine individuals could not be analyzed due to technical issues. Additionally, DNA was extracted from blood samples to account for genetic influences in the analysis. The z-GIG study was approved by the Zurich Cantonal Ethics Committee (KEK, BASEC No. 2017-02083), and all participants provided written informed consent.

### Design and Bias Adjustment

The study draws on several methodological strategies: (1) longitudinal PV measures were used, with newly collected experimental and questionnaire data minimizing common method variances (Gini & Pozzoli, 2013; Rutter et al., 2001); (2) genetically informed inverse probability weighting (IPW), incorporating psychosocial baseline confounders, polygenic risk scores, and genetic ancestry was applied to mitigate bias from both selection into treatment (i.e., PV) and selection into the enriched sample (Hernan, 2006; D. B. Rubin, 1997) and to enable covariate adjustment without reducing degrees of freedom in the outcome model (Rosenbaum & Rubin, 1984); (3) missing data were imputed using a random forest method, minimizing bias from list-wise deletion (D. B. Rubin, 1976); (4) confounding social adversities such as PV perpetration, sexual victimization, and childhood maltreatment (corporal punishment), were accounted for (Baldwin et al., 2023; Catone et al., 2015; Geoffroy et al., 2018; Stapinski et al., 2014); and (5) longitudinal data were leveraged by adjusting for key time-varying confounders during the victimization period (Zuber et al., 2023), including substance use, psychological help-seeking, and exercise, with principal components (explaining 75% of variance) used to maximize statistical power; and (6) since depression is also associated with biased attention (Craske et al., 2009) cumulative depression was included as covariate, averaged across five waves using the SBQ internalizing scale (Tremblay, 1994; z-proso Project Team, 2024), as well as subjective stress at age 20. References for additional scales are provided in Supporting Table S1.

### Experimental Paradigms and Surveys

Participants first completed a free-viewing task (Lisk et al., 2020; Shechner et al., 2012), followed by the AIHQ questionnaire, and finally a morphing task (Ito et al., 2017). Facial expressions for the tasks were selected from the Chicago Face Database (Ma et al., 2015), comprising 42 angry–happy image pairs (21 male, 21 female) with closed-mouth expressions, colored on a white background, and matched to the participants’ age group, with predominantly European-like features for consistency with the sample. Stimulus luminance (mean = 0.814; SD = 0.030) was comparable across expressions (*t* (_82_) = –1.185; *p* = 0.240), with any residual differences accounted for in analyses. The viewing distance was about 1 m, with a screen resolution of 1280 × 1024 px. Experiments were conducted in a room with constant artificial lighting, without daylight.

#### Free-viewing?

Each of the 42 faces was presented with a happy and an angry expression, randomly positioned on the left or right (537 × 378 px each) against a 50% gray background. Visual stimuli were displayed on a 19-inch Philips 109S2 CRT monitor (1280 × 1024 px resolution, 36.5 cm screen width, 4:3 aspect ratio, and 85 Hz refresh rate). The order of face pairs was randomized once and presented uniformly to all participants for consistency. Eye gaze (*x, y* coordinates) was recorded using an EyeLink 1000 (v4.51) at 500 Hz (monocular desktop mounted, chin-stabilized setup; 1280 × 1024 px resolution). Each trial began with a 2 s fixation on a black dot centered on the background. To maintain their attention, participants were informed they would perform a dot-probe task (MacLeod et al., 1986); after 5,000 ms, the faces were replaced by a randomly positioned pair of 20 px black dots (left or right, arranged either horizontally or vertically, with a center-to-center distance of 30 px). Participants responded as quickly as possible by pressing the left arrow key for horizontally arranged dots and the up arrow key for vertical dots. After two test trials, gaze coordinates and reaction times were recorded. Ten participants, including two victims, were excluded from the eye-tracking analysis due to poor calibration, excessive eye movement, or gaze instability (e.g., caused by glasses or poor concentration). Attention in the free-viewing task was assessed using adapted measures previously described (Stilling et al., 2018): (1) attentional maintenance, defined as the ratio of fixations on angry versus happy faces; (2) initial orienting, defined as probability whether the first fixation was directed toward the angry face; (3) disengagement, operationalized as the time elapsed before the initial fixation was broken; and (4) hyperscanning, indexed through comparisons of scan path lengths (in px). Square regions of interest (247 × 247 px) were defined to encompass the entire facial area of each stimulus.

#### Morphing

For each of the 42 faces, 101 morphed frames were generated (Ito et al., 2017), ranging from happy to angry expressions. Participants used an interactive morphing interface to adjust a static face (896 × 630 px, 50% gray background) along this continuum, selecting their perceived neutral point between happy and angry as quickly as possible by moving the computer mouse and confirming their choice with the space bar. Starting frames were randomly selected. A black fixation dot was presented for 2 s at the center of the screen between trials. After two test trials, mouse endpoint coordinates and reaction times were recorded.

#### AIHQ

The AIHQ (Combs et al., 2007) measured hostile social-cognitive biases in response to ambiguous negative situations varying in intentionality. It includes the key indices hostility bias (hostile attribution) and aggression bias (reactive aggression) among others. In the z-proso panel at age 9, participants completed a similar instrument (Dodge & Coie, 1987; Dodge et al., 1990), which closely parallels the AIHQ, except for the absence of a blame score (therefore not included in this analysis), and could be considered its child version. To ensure compatibility, two raters scored responses at ages 9 and 21 with the AIHQ scale (Krippendorff’s *α*: mean = 0.768; SD = 0.096). All scores were normalized.

### Genetic Data

DNA of 153 (84.1%) participants was isolated from EDTA blood at LIFE & BRAIN GmbH (Bonn, Germany) for high-throughput bulk analysis and genotyped according to the protocol previously described (Jagannath et al., 2020) using the Illumina Infinium GSA array, mapped to GRCh38. Polygenic risk scores, used in IPW, were calculated from European GWAS summary statistics for relevant psychiatric traits [attention-deficit/hyperactivity disorder (Demontis et al., 2023), generalized anxiety disorder (Otowa et al., 2016), major depressive disorder (Wray et al., 2018), and panic disorder (Forstner et al., 2021)] converted to GRCh38 using the UCSC LiftOver Tool; quality control followed established guidelines (Choi et al., 2020). The first four principal components of genetic ancestry (Supporting Fig. S3) were calculated to account for population structure (Price et al., 2006). PLINK 1.90b7 (Purcell et al., 2007) and GNU awk 5.1.0 was used for DNA bioinformatics.

### Statistical Analysis

Analyses were performed in R 4.4.3 using custom scripts. IPW was applied, incorporating baseline biopsychosocial covariates, psychiatric polygenic risk scores, and genetic ancestry. Missing data were imputed using missRanger v2.6.0 (Stekhoven & Buhlmann, 2012). The larger control group improved covariate balance and comparability with the PV group. Propensity score diagnostics indicated successful covariate balance (for diagnostics see the Supporting Information).

For the free-viewing and morphing tasks, IPW-weighted mixed models with a random intercept structure were estimated to account for subject- and trial-specific effects. For the binary outcome (initial orienting toward angry faces), a linear probability mixed model was used due to its straight-forward interpretability; effect estimates and significance patterns showed no meaningful differences compared to binomial models. PV effects on AIHQ scores were estimated using IPW-weighted mixed-effects models with two time-points. All standard errors were estimated using 100,000 bootstrap replications to account for pseudo-population variance introduced by IPW.

## Results

Sample characteristics in the weighted dataset were well balanced between the PV and non-PV control group (Table 1).

**Table 1.**
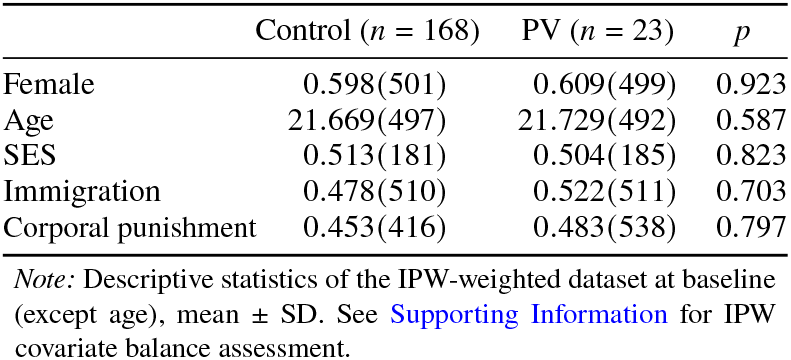
Descriptive statistics of the weighted dataset.

|  | Control (*n* = 168) | PV (*n* = 23) | *p* |
| --- | --- | --- | --- |
| Female | 0.598(501) | 0.609(499) | 0.923 |
| Age | 21.669(497) | 21.729(492) | 0.587 |
| SES | 0.513(181) | 0.504(185) | 0.823 |
| Immigration | 0.478(510) | 0.522(511) | 0.703 |
| Corporal punishment | 0.453(416) | 0.483(538) | 0.797 |

*Note:* Descriptive statistics of the IPW-weighted dataset at baseline (except age), mean ± SD. See Supporting Information for IPW covariate balance assessment.

### Free-viewing

Victims spent less time fixating on angry faces than controls when initially orienting to the happy stimulus; this difference was attenuated when first orienting to angry faces (Table 2A). No significant group differences were found in the probability of initially orienting toward either stimulus (Table 2B). When initially orienting to angry faces, victims disengaged faster than controls (Table 2C), with no significant group difference observed for disengagement from happy faces. Scan paths were about 20% longer in victims than controls (Table 2D). As expected in a free-viewing design with prolonged exposure, reaction times did not differ by group (*β* = 0.009; 95% CI = [–0.005, 0.023]; *p* = 0.231).

**Table 2.** Results for the free-viewing task (n = 179).

A. Attentional maintenance on angry stimulus (ratio)
| | $\beta$ | 95% CI | *p* |
| --- | --- | --- | --- |
| (Intercept) | 1.194 | [0.995, 1.392] | <0.001 |
| PV | -0.244 | [-0.433, -0.056] | 0.011 |
| Angry First | 0.192 | [0.101, 0.283] | <0.001 |
| PV × Angry First | 0.145 | [0.008, 0.281] | 0.038 |

B. Initial orienting to angry stimulus (probability)
| | $\beta$ | 95% CI | *p* |
| --- | --- | --- | --- |
| (Intercept) | 0.444 | [-0.365, 0.521] | <0.001 |
| PV | 0.002 | [-0.063, 0.068] | 0.949 |

C. Disengagement from initial stimulus (ms)
| | $\beta$ | 95% CI | *p* |
| --- | --- | --- | --- |
| (Intercept) | 435.502 | [334.570, 535.570] | <0.001 |
| PV | -54.992 | [-153.830, 43.991] | 0.276 |
| Angry First | 40.558 | [10.567, 70.433] | 0.008 |
| PV × Angry First | -45.300 | [-83.891, -6.827] | 0.021 |

D. Hyperscanning (log scan path in pixels)
| | $\beta$ | 95% CI | *p* |
| --- | --- | --- | --- |
| (Intercept) | 8.889 | [7.714, 9.066] | <0.001 |
| PV | 0.186 | [0.007, 0.365] | 0.042 |

*Note:* IPW mixed models. Bootstrapped standard errors (100K reps), BCa confidence intervals. PV = peer victimization; Angry First = first fixation on angry face. Covariates included but not shown.

### AIHQ

In both groups, hostile attribution bias and reactive aggression decreased over time. However, victims followed a distinct trajectory. Compared to controls, they showed a weaker decline in hostile attribution bias and a stronger decline in reactive aggression (Table 3; Fig. 1).

**Table 3.**
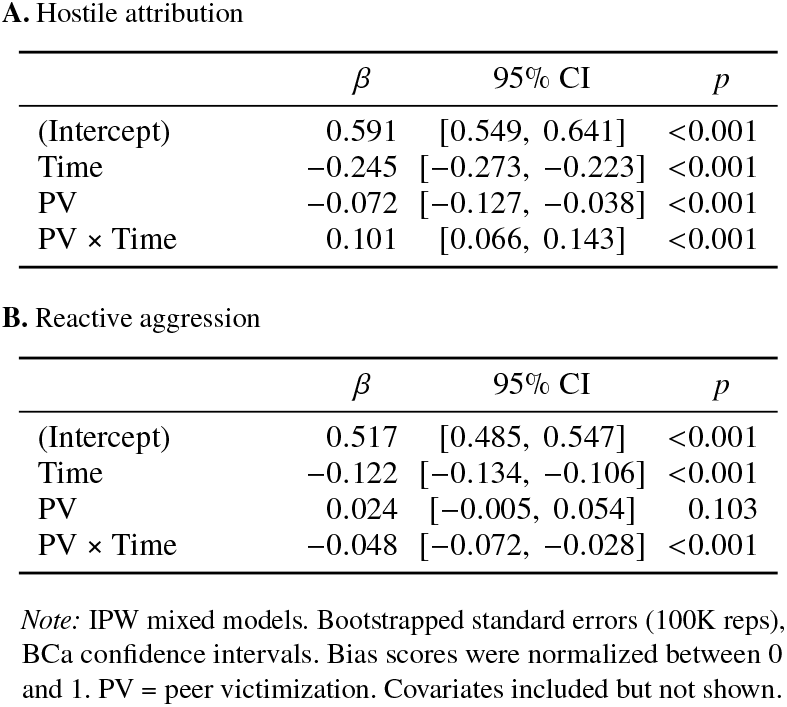
Results from the AIHQ (*n* = 191).

**Figure 1.**
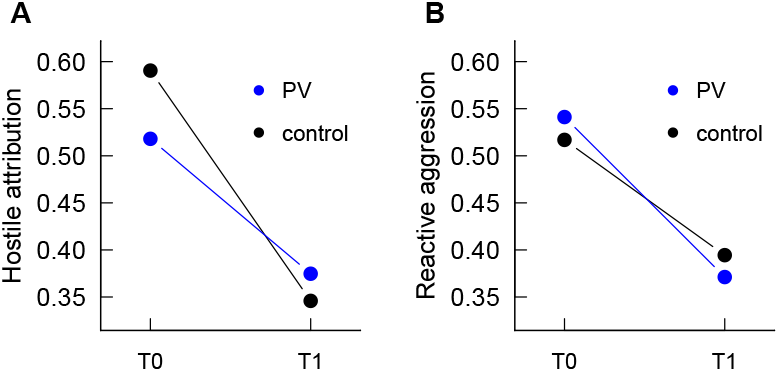
Estimated marginal means of social-cognitive biases. T0 = 9y, T1 = 22y. Biases decreased over time in both groups, but with diverging trajectories. **(A)** Hostile attribution was initially lower in victims but surpassed that of controls at follow-up. **(B)** Reactive aggression started slightly higher in victims but declined more steeply, resulting in lower levels than controls.

### Morphing

In the morphing task, victims placed their neutral point further toward the angry end of the morph continuum compared to controls (Table 4). Reaction times for victims did not differ significantly between groups (*β* = −0.032; 95% CI = [−0.099, 0.034]; *p* = 0.347).

**Table 4.** Results from the morphing task (*n* = 191).

| | $\beta$ | 95% CI | $p$ |
| --- | --- | --- | --- |
| (Intercept) | 50.110 | [45.979, 54.273] | <0.001 |
| PV | 5.684 | [2.139, 9.157] | 0.002 |

Note: IPW mixed model. Bootstrapped standard errors (100K reps), BCA confidence intervals. Scores reflect the frame number (happy = 1, angry = 100) at which faces were judged as aggressive. PV = peer victimization. Covariates included but not shown.

## Discussion

Evidence from this study links adolescent peer victimization (PV) to disruptions in normative psychosocial development and impaired social functioning in young adulthood. Using a counterfactual approach, analyses showed that, compared to a non-PV control group, young adults previously exposed to severe adolescent PV (victims) showed vigilance-avoidance-driven attentional shifts, an interpretational bias toward perceiving angrier faces as neutral, and—over the course of adolescence—kept showing more hostile attribution and less reactive aggression than controls.

Reduced attentional maintenance on angry faces among victims compared to controls aligns with findings in physically abused children, who tend to divert their attention away from threat—a behavior linked to social phobia and PTSD (Y. Chen et al., 2002; Jaffee, 2017; Pine et al., 2005). It may also reflect a broader tendency to avoid eye contact with angry—and thus likely more dominant—others, potentially indicating a subordinate social stance (Sapolsky, 2004). Victims’ initial orienting to angry faces appeared to enhance attentional engagement among victims; disengagement from the first fixation was attenuated—i.e., modestly slower—followed by greater cumulative attentional maintenance. Alongside the observed one-fifth longer scan paths, this pattern suggests a sequence of early disengagement followed by repeated returns to threat-related stimuli. Such a profile may reflect a strategic attentional response to social threat (Gü nther et al., 2021; Kircanski et al., 2015; Rapee et al., 2023), in which initial avoidance is followed by controlled re-engagement aimed at mitigating emotional impact—a pattern consistent with vigilance–avoidance dynamics (Hoepfel et al., 2022).

While such avoidance strategies may buffer stress in the short term, this selective attention profile observed among victims—resembling patterns described in social anxiety disorder and PTSD (N. T. M. Chen & Clarke, 2017; Wald et al., 2011; Williams et al., 2024)—may reflect a seemingly resilient yet dissociative adaptation during adolescence (Herzog et al., 2018). Over time, this profile may impose sustained cognitive strain (Gü nther et al., 2021; Ke et al., 2022; McLaughlin et al., 2015), potentially contributing to impaired inhibitory control and cognitive decline in adulthood (Avramescu et al., 2024; Pantoja-Urbán et al., 2024), underscoring the relevance of integrative neurobiological and psychological approaches to understanding developmental stress responses.

Interestingly, first fixation–related patterns emerged, even though groups did not differ in the likelihood of initially fixating on a particular emotion. Effects specific to angry faces may mirror those reported in a recent meta-analysis (Gü nther et al., 2021), yet the modulation by first fixation raises open questions regarding the role of first fixations in emotion processing, which should be explored in more targeted paradigms.

Longitudinal estimates of social-cognitive biases suggest that PV is associated with diverging developmental trajectories. Victims’ flatter decrease in hostile attribution from childhood to adolescence suggests that PV may interfere with the typical normative decline in perceived hostility. This finding complements previous studies showing that victimized adolescents perceive ambiguous social situations as more threatening (Schacter et al., 2024; Ziv et al., 2013) with the finding that PV may play a role in the developmental progression of such cognitive biases. In contrast, the steeper decline in reactive aggression among victims suggests a more pronounced reduction in their tendency to endorse aggressive responses over time. Interestingly, animal models have similarly shown that repeated social stress in adolescence is associated with reduced aggression as well as increased avoidance, mediated by reduced HPA axis activity, highlighting adolescence as a sensitive period (Ver Hoeve et al., 2013). While previous research has linked PV to elevated reactive aggression—primarily based on cross-sectional designs (Card & Little, 2006)—our findings may complement this literature by capturing a blunting of affective reactivity within victims over time.

Consequently, bullies may not only select victims who are less likely to retaliate effectively (Volk et al., 2022), but may also contribute to a maladaptive socialization process that increasingly disempowers victims, diminishing their capacity to respond assertively or redirect frustration. This aligns with findings that individuals of lower social status exhibit reduced displacement aggression and possess fewer means to buffer or cope with frustration, such as social affiliation or support networks, e.g., with fellow subordinate individuals (Aupperle et al., 2012; Chen Zeng et al., 2022; Glowacki et al., 2020; Kruglanski et al., 2023; Sapolsky, 2004). This challenges prevailing theories emphasizing retaliation (Camodeca & Goossens, 2005; Herd & Kim-Spoon, 2021) and highlights the importance of the present longitudinal findings.

In a broader sense, such patterns may contribute to the friendship and romantic barriers frequently reported by victims (Arseneault, 2018; D’Urso et al., 2023; Kochel et al., 2012), as general social approach behavior may also be adversely affected. Indeed, victims often report fewer sexual partners (Gallup et al., 2009; Volk et al., 2022), a pattern that may reflect reproductive suppression as commonly observed in subordinate individuals in humans and other animals (Sapolsky, 2004). Anger suppression was also observed in depressive individuals in line with evidence suggesting a negative association between depression and externalized aggression (Riley et al., 1989).

One potential mechanism may involve elevated cortisol levels among victims, consistent with evidence linking higher cortisol to reduced reactive aggression (Pfattheicher & Keller, 2014). Notably, victims exhibited lower hostile attribution in childhood compared to controls, suggesting a more trusting or socially naïve interpretive style. This reduced sensitivity to potential interpersonal threat may have increased their vulnerability to PV.

Victims exhibited a more anger-shifted neutral point on the happy–angry continuum relative to controls, suggesting a cognitive bias in facial emotion processing. One possible explanation is reduced perceptual sensitivity to angry expressions—potentially reflecting habituation—consistent with prior findings that acute stress can dampen anger perception in healthy children (F. S. Chen et al., 2014), a pattern similarly reported in victims of terrorist attacks (Wormwood et al., 2016). Emotional numbing of this kind is commonly observed in PTSD (Plana et al., 2014), suggesting that PV-induced trauma effects (e.g., Ossa et al., 2019) might outweigh those associated with social anxiety. Thus, while research on child maltreatment has often highlighted hypersensitivity to threat (e.g., Gibb et al., 2009; Pollak and Kistler, 2002), our findings in youth instead point toward blunted threat sensitivity as a possible downstream effect—consistent with desensitization hypotheses (Kellij et al., 2022).

The underlying mechanism could be a reduced neuronal responsivity to negative emotional stimuli. Lesion studies suggest that the ventromedial part of the prefrontal cortex (PFC) and orbitofrontal regions play a crucial role in recognizing facial emotions, especially subtle emotional cues and signs of anger (Davidson et al., 2000; Heberlein et al., 2008; Hiser & Koenigs, 2018; Tsuchida & Fellows, 2012). Supporting this, fNIRS studies have shown reduced oxy-Hb responses in temporal regions during negative face processing in ADHD, a disorder similarly characterized by difficulties in facial emotion recognition (Ichikawa et al., 2014). A similar bias toward perceiving happiness in ambiguous or negative facial expressions has been observed alongside reduced aggression (Penton-Voak et al., 2013) and in individuals with amygdala distortions (Monk et al., 2006; Sato et al., 2002), suggesting a broader neural basis involving vmPFC, OFC, amygdala, and temporal circuits.

Alternatively, the pattern may reflect increased top-down modulation during emotion interpretation (McLaughlin et al., 2015). Socially anxious individuals interpret happy faces more slowly than angry ones (Maoz et al., 2016), suggesting greater cognitive load when processing positive expressions. Similarly, in children exposed to family violence, the typically negative correlation between amygdala and PFC activity was reversed, indicating heightened self-regulatory engagement in response to emotional stimuli (Taylor et al., 2006). In fact, victims in our study may have selected a facial expression with higher anger intensity as neutral in order to reduce ambiguity and cognitive demand. This interpretation is consistent with the time pressure imposed by the task and the absence of significant group differences in reaction times. Yet another explanation is that victims, being arguably more experienced with anger cues, may be more accurate in identifying anger—suggesting that the actual bias lies with the control group, who tend to flag anger too readily.

### Limitations

Although based on longitudinal data, the study analyzed PV as a binary outcome; youth who were always in the highest decile with victimization but never with perpetration between the ages of 11 and 20 were classified victims. This measurement strategy has arguably identified a distinctive group and provided valuable insights. It could, however, be seen as a limitation in that more detailed statistical analysis of assumed PV patterns (e.g. early vs. late, acute vs. chronic) was not possible; future research may analyze larger samples in which the resulting PV clusters reach a sufficient size for statistical analysis. Moreover, the lack of information about self-perception as a victim made it necessary to ascribe the victim role using observer-defined criteria which may cause underestimation of PV effects.

In the free-viewing task, the anxiety level was not modified so it remains unclear whether victims have a different threshold depending on increasing emotional levels (Cisler & Koster, 2010; Grossheinrich et al., 2022). Also, because socially anxious individuals may perceive emotional expressions—including happy faces—as threatening (Liu et al., 2021), the happy–angry contrast may limit the interpretability of underlying attentional mechanisms.

## Conclusion

Together, findings suggest that adolescent PV may initiate developmental cascades affecting attentional and social-cognitive functioning, as reflected in disrupted attention to threat, biased attribution of intent, blunted affective re-activity, and impaired facial emotion processing—all of which were observed in young adulthood. The observed patterns among PV victims resemble those seen in social anxiety disorder and post-traumatic stress disorder, which may shed light on the mechanisms behind social withdrawal and loneliness frequently reported by victims. These insights may help inform therapeutic approaches aimed at mitigating victims’ social difficulties. Finally, the enduring psychosocial ramifications of adolescent PV highlight its relevance as a public health concern and call for more targeted, youth-focused research.

## Acknowledgments

This work was supported by funding from the Swiss National Science Foundation (Grants 10531C-197964 to MJS, 405240-69025, 100013 116829, 100014 132124, 100014 - 149979, 10FI14 170409), the Jacobs Foundation (Grants 2010-888, 2013-1081-1), the Jacobs Center for Productive Youth Development, the Swiss Federal Office of Public Health (Grants 2.001391, 8.000665), the Canton of Zurich’s Department of Education, the Swiss Federal Commission on Migration (Grants 03-901 (IMES), E-05-1076), the Julius Baer Foundation, and the Visana Foundation.

We gratefully acknowledge all individuals from the z-proso and its z-GIG subsample for their voluntary participation. During the work on this paper, Jens Heumann was a fellow of the International Max Planck Research School on the Life Course (LIFE; https://www.imprs-life.mpg.de/).

## SUPPORTING INFORMATION

### Peer victimization variable definition

PV classification was based on observer-defined criteria. This was necessary because, in the z-proso questionnaires, victims were only asked about experienced and exerted peer adversity but not about self-perception as a victim. Classification required consideration of subordinate experiences (sometimes already referred to as “victimization”) as well as dominant behavior (sometimes referred to as “perpetration”); if both occurred, the case was considered as dynamic behavior rather than victimization, due to an assumed difference in situational appraisal (Kemeny, 2003). Further, assumptions had to be made about what adversity may have occurred during the gap years between waves, since no assessment was conducted then (Fig. S1). Also, to define an unaffected control group represented in model intercepts, it had to be taken into account that participants may also have faced other impactful adversities, such as sexual adversity—which was not consistently measured—and online and dating adversity—which were not measured at all. Finally, declining trends in adversity severity across the decade of measurement had to be considered.

**Figure S1.**
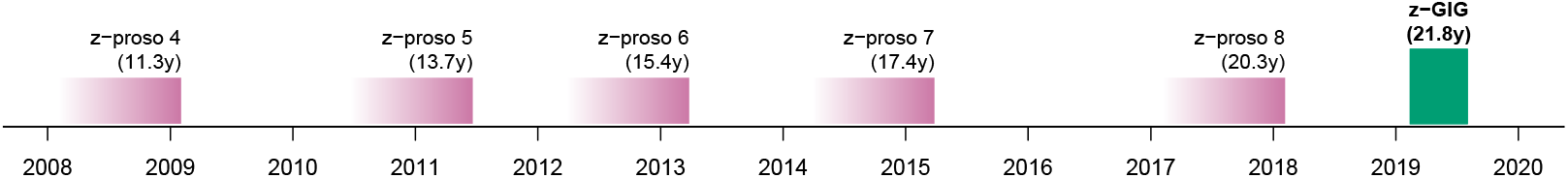
Study timeline showing the schedule of peer adversity assessments in the z-proso panel. Each bar’s right edge corresponds to a survey wave, while the leftward gradient represents the 12-month retrospective reporting period. Gaps indicate periods when no data were collected.

To implement this classification, the following approach was adopted to classify victims of peer adversity: (1) consistently measured incidence across the primary adversity domains—verbal, social, and physical aggression—was assessed using Likert scales (1 = never, 2 = 1 time, 3 = 3 times, 4 = about once a month, 5 = about once a week, 6 = about every day) and summed and normalized separately for experienced and exerted adversity; (2) severe occurrence at each measurement point was defined as scoring above the 90th percentile (subordinate experiences: 2009 = 0.36, 2011 = 0.36, 2013 = 0.28, 2015 = 0.20, 2018 = 0.20; dominant behavior: 2009 = 0.24, 2011 = 0.36, 2013 = 0.28, 2015 = 0.20, 2018 = 0.16), capturing the downward trend; (3) after subtracting dominant behavior, 23 participants with positive scores only in subordinate experiences were coded as victims in the dummy variable and 168 as controls. Controls consisted of 90 individuals (53.6%) with zero primary adversity, 32 (19.0%) with dominant behavior, and 46 (27.4%) with both subordinate experiences and dominant behavior. These behaviors, along with known sexual adversity, were statistically controlled in the model to ensure that the control group (represented in the intercept) was not confounded by other forms of adversity.

**Figure S2.**
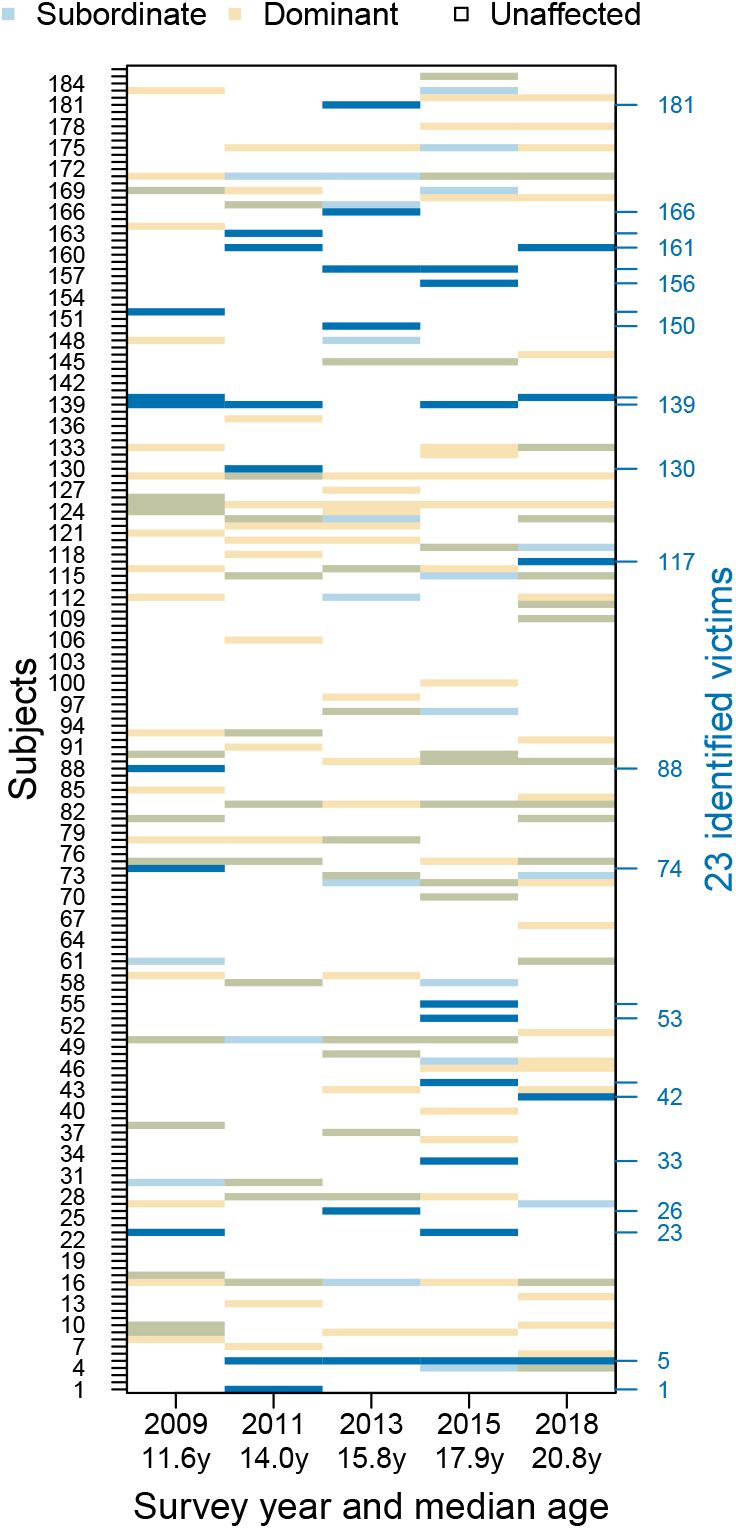
Longitudinal patterns of peer adversity. Each row corresponds to one participant, with subordinate experiences (blue) and dominant experiences (orange) displayed across survey waves spanning ages 11 to 20. Individuals with repeated subordinate experiences and no dominant behavior were classified as peer-victimized (*n* = 23), comprising the victim group at age 22.

### Inverse probability weighting (IPW)

Across all analyses, IPW (Hernán & Robins, 2020) was applied to account for both treatment (PV) selection bias and stratified sampling. This counterfactual approach generates a pseudo-population approximating a randomized trial and aids causal interpretation and is generally recommended for observational studies, especially in PV research (Baldwin et al., 2023). To further reduce confounding, weights were genetically informed by psychiatric polygenic risk scores (PRSs), genetic ancestry, and biopsychosocial baseline confounders. Confounders were defined as influencing both PV and outcome (Rubin, 1997). Weights for estimating the average treatment effect on the treated (ATT, Eq. 1) were derived from propensity scores for PV calculated via a logit model,

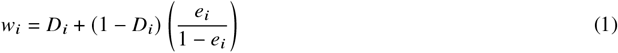

where *D* denotes the PV dummy, and *e* the propensity score (Morgan & Winship, 2015). The ATT was chosen as actual victims represent the primary population of interest. To reduce the influence of extreme values, weights were trimmed at the 5th and 95th percentiles. Baseline data for the propensity score model were taken from: (i) self-reports (adolescents, teachers, and parents) from the first three z-proso waves (ages 7–9) prior to the measurement of PV; (ii) PRS for attention-deficit/hyperactivity disorder (ADHD), generalized anxiety disorder (GAD), major depressive disorder (MD), and panic disorder (PD); and (iii) first four principal components from genotyping based on eigenvalue analysis, which showed a clear inflection point at the fourth component with an eigenvalue near 1 (Fig. S3) (Cattell, 1966; Price et al., 2006) to account for population structure within the sample. Missing values (0.5–6% across all variables) were imputed using random forests [missRanger 2.6.0; (Stekhoven & Buhlmann, 2012)]. A full list of confounders and corresponding references of expert literature is provided in Table S1.

**Figure S3.**
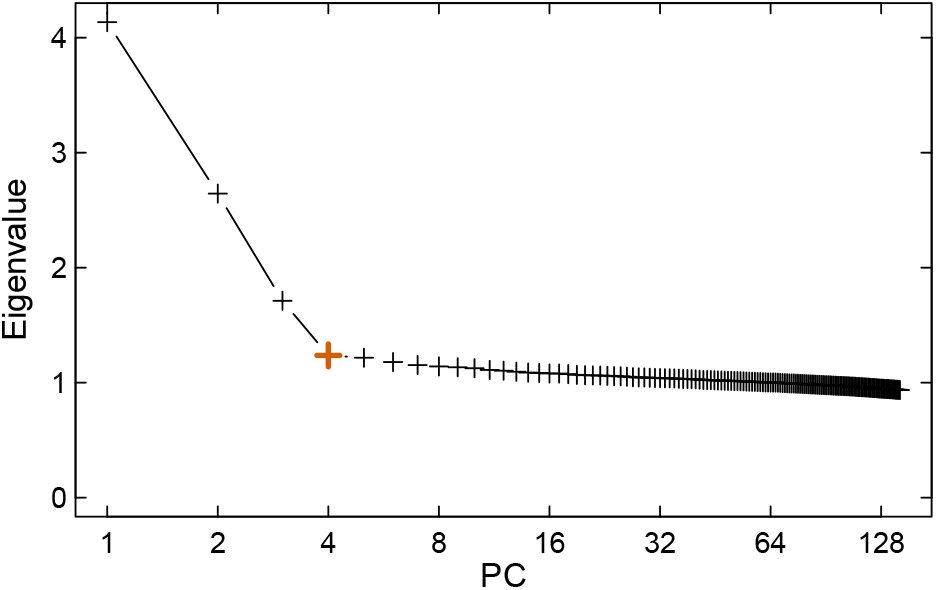
Scree plot of eigenvalues from genotyping data. An elbow at the fourth component suggests that the first four dimensions capture ancestry-related structure.

To minimize extrapolation bias, ATT estimation was restricted to victims with propensity scores within the range of the controls (common support) and excluded two of the original 25 victims due to lack of appropriate matches (Austin & Stuart, 2017; Stuart, 2010), resulting in the final group of 23 victims. IPW weights *w* were applied as regression weights throughout the analyses using bootstrap standard errors with 100,000 replications and bias-corrected and accelerated confidence intervals (BCa) to account for the pseudo-population variance (Davison & Hinkley, 1997).

### IPW diagnostics

The maximum absolute standardized difference was 0.198 (Fig. S4A). Mean of expected weights (0.259) and estimated weights (0.250) differed by -0.009, indicating good agreement and stability (Reifeis & Hudgens, 2022). Propensity scores of victims and controls demonstrated sufficient overlap (Fig. S4B).

**Figure S4.**
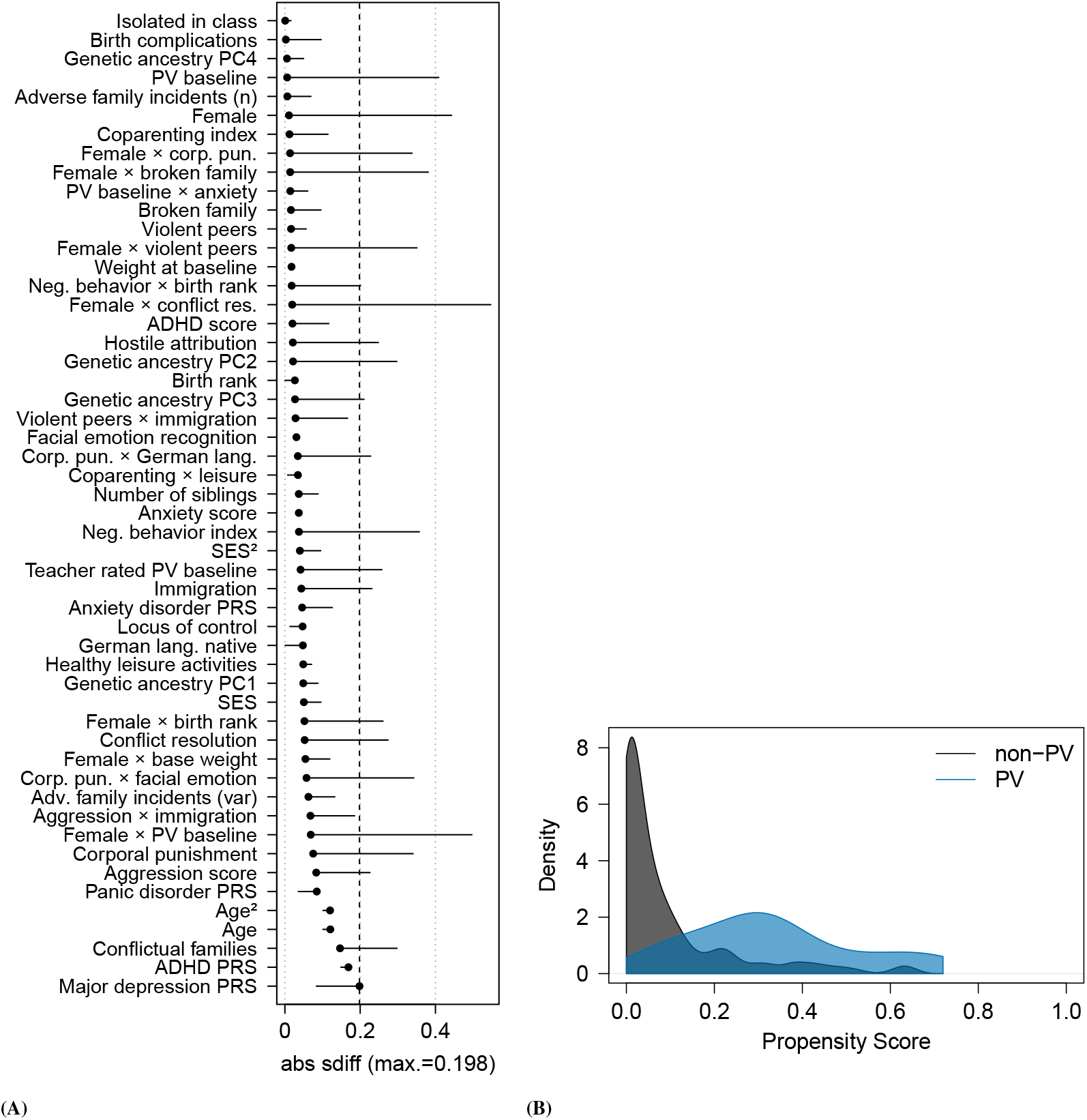
(**A**) Covariate balance before and after IPW. Absolute standardized differences (ASD) are shown for each covariate used in weighting. Vertical dashed line marks the maximum ASD after weighting (0.198), indicating successful covariate balancing. (**B**) Propensity score distributions. Histogram of propensity scores for treated and control groups in the final analytic sample, indicating sufficient overlap for group comparability.

**Table S1.** Confounders relevant to the relationship between PV and health outcomes, identified through expert knowledge and literature.

| Confounder |
| --- |
| ADHD-score, conflict resolution, corporal punishment<sup>a</sup>, isolated in class, teacher rated bullying (Holt & Espelage, 2007; Lereya et al., 2015; Zych et al., 2020) |
| Adverse family incidents, broken family, conflictual family (Forster et al., 2020; Lereya et al., 2013) |
| Age (Eastman et al., 2018) |
| Aggression, anxiety<sup>b</sup> (Espelage et al., 2001; Schoeler et al., 2018) |
| Birth complications (DiLalla et al., 2021) |
| Birth rank (Dantchev & Wolke, 2019) |
| Coparenting<sup>c</sup> (Martínez et al., 2019) |
| Depression<sup>b</sup> (Penner-Goeke et al., 2023; Rose et al., 2016) |
| Facial emotion recognition<sup>d</sup> (Dede et al., 2021) |
| Genetic disposition<sup>e</sup> (Brendgen et al., 2015) |
| Healthy leisure activities<sup>f</sup>, body weight (at baseline) (Puhl et al., 2011) |
| Hostile attribution (Perhamus et al., 2022) |
| Immigration, German language native (Qin et al., 2008) |
| Negative behavior in childhood<sup>f</sup>, self-control (Cook et al., 2010; Jennings et al., 2011; Robson et al., 2020; Zych et al., 2020) |
| Number of siblings (Karišik, 2019; Wu et al., 2023) |
| Sex (Casper & Card, 2017; Schoeler et al., 2018; White & Kaffman, 2020) |
| Socio-economic status<sup>g</sup> [SES; (Delprato et al., 2017; Shanahan et al., 2022; Tippet & Wolke, 2014; Wong & Schonlau, 2013)] |
| Violent peers (Duffy & Nesdale, 2009) |

Notes: <sup>a</sup>Corporal punishment was assessed using the Social Behavior Questionnaire (SBQ; Murray et al., 2019). <sup>b</sup>Polygenic risk scores (PRS) were used for both anxiety and depression, alongside an observed anxiety score. <sup>c</sup>The coparenting score—derived from items of the Alabama Parenting Questionnaire (APQ; Murray et al., 2021)—reflected the degree of agreement between father and mother in their survey responses on parenting. <sup>d</sup>Accuracy in facial emotion recognition was scored using correct responses on the Assessing Children's Emotional Skills (ACES; Schultz et al., 2004) instrument, administered as part of z-proso. <sup>e</sup>Genetic predispositions were measured using PRS derived from relevant GWAS datasets. <sup>f</sup>Healthy leisure and negative behavior scores were constructed from respective z-proso survey items. <sup>g</sup>Socio-economic status (SES) was calculated based on parental income, education, and occupational prestige (ISEI). Additional scale references can be found in the z-proso project handbook (z-proso Project Team, 2024).

## References

Armstrong, T., & Olatunji, B. O. (2012). Eye tracking of attention in the affective disorders: A meta-analytic review and synthesis. Clinical Psychology Review, 32(8), 704–723. 10.1016/j.cpr.2012.09.004.

Arseneault, L. (2018). Annual Research Review: The persistent and pervasive impact of being bullied in childhood and adolescence: Implications for policy and practice. Journal of Child Psychology and Psychiatry, 59(4), 405–421. 10.1111/jcpp.12841.

Aupperle, R. L., Melrose, A. J., Stein, M. B., & Paulus, M. P. (2012). Executive function and PTSD: Disengaging from trauma. Neuropharmacology, 62(2), 686–694. 10.1016/j.neuropharm.2011.02.008.

Avramescu, R. G., Capolicchio, T., & Flores, C. (2024). Dynamic Insights into Dopamine Axon Growth in Adolescence and its Implications for Psychiatric Risk. Current Opinion in Behavioral Sciences, 59, 101435. 10.1016/j.cobeha.2024.101435.

Baldwin, J. R., Wang, B., Karwatowska, L., Schoeler, T., Tsaligopoulou, A., Munafò, M.R., & Pingault, J.-B. (2023). Childhood Maltreatment and Mental Health Problems: A Systematic Review and Meta-Analysis of Quasi-Experimental Studies. American Journal of Psychiatry, 180(2), 117–126. 10.1176/appi.ajp.20220174.

Bishop, S. J. (2007). Neurocognitive mechanisms of anxiety: An integrative account. Trends in Cognitive Sciences, 11(7), 307–316. 10.1016/j.tics.2007.05.008.

Biswas, T., Scott, J. G., Munir, K., Thomas, H. J., Huda, M. M., Hasan, M. M., David De Vries, T., Baxter, J., & Mamun, A. A. (2020). Global variation in the prevalence of bullying victimisation amongst adolescents: Role of peer and parental supports. EClinicalMedicine, 20, 100276. 10.1016/j.eclinm.2020.100276.

Briggs-Gowan, M. J., Pollak, S. D., Grasso, D., Voss, J., Mian, N. D., Zobel, E., McCarthy, K. J., Wakschlag, L. S., & Pine, D. S. (2015). Attention bias and anxiety in young children exposed to family violence. Journal of Child Psychology and Psychiatry, 56(11), 1194– 1201. 10.1111/jcpp.12397.

Buss, D. M., & Shackelford, T. K. (1997). Human aggression in evolutionary psychological perspective. Clinical Psychology Review, 17(6), 605–619. 10.1016/S0272-7358(97)00037-8.

Camodeca, M., & Goossens, F. A. (2005). Aggression, social cognitions, anger and sadness in bullies and victims. Journal of Child Psychology and Psychiatry, 46(2), 186–197. 10.1111/j.14697610.2004.00347.x.

Card, N. A., & Little, T. D. (2006). Proactive and reactive aggression in childhood and adolescence: A meta-analysis of differential relations with psychosocial adjustment. International Journal of Behavioral Development, 30(5), 466–480. 10.1177/0165025406071904.

Catone, G., Marwaha, S., Kuipers, E., Lennox, B., Freeman, D., Bebbington, P., & Broome, M. (2015). Bullying victimisation and risk of psychotic phenomena: Analyses of British national survey data. The Lancet Psychiatry, 2(7), 618–624. 10.1016/S2215-0366(15)00055-3.

Ceccarini, F., Colpizzi, I., & Caudek, C. (2024). Age-dependent changes in the anger superiority effect: Evidence from a visual search task. Psychonomic Bulletin & Review, 31(4), 1704–1713. 10.3758/s13423-023-02401-3.

Chen, F. S., Schmitz, J., Domes, G., Tuschen-Caffier, B., & Heinrichs, M. (2014). Effects of acute social stress on emotion processing in children. Psychoneuroendocrinology, 40, 91–95. 10.1016/j.psyneuen.2013.11.003.

Chen, N. T. M., & Clarke, P. J. F. (2017). Gaze-Based Assessments of Vigilance and Avoidance in Social Anxiety: A Review. Current Psychiatry Reports, 19(9), 59. 10.1007/s11920-017-0808-4.

Chen, Y., Ehlers, A., Clark, D., & Mansell, W. (2002). Patients with generalized social phobia direct their attention away from faces. Behaviour Research and Therapy, 40(6), 677–687. 10.1016/S0005-7967(01)00086-9.

Chen Zeng, T., Cheng, J. T., & Henrich, J. (2022). Dominance in humans. Philosophical Transactions of the Royal Society B: Biological Sciences, 377(1845), 20200451. 10.1098/rstb.2020.0451.

Choi, S. W., Mak, T. S.-H., & O’Reilly, P. F. (2020). Tutorial: A guide to performing polygenic risk score analyses. Nature Protocols, 15(9), 2759–2772. 10.1038/s41596-020-0353-1.

Cisler, J. M., & Koster, E. H. (2010). Mechanisms of attentional biases towards threat in anxiety disorders: An integrative review. Clinical Psychology Review, 30(2), 203–216. 10.1016/j.cpr.2009.11.003.

Claudino, R. G. E., De Lima, L. K. S., De Assis, E. D. B., & Torro, N. (2019). Facial expressions and eye tracking in individuals with social anxiety disorder: A systematic review. Psicologia: Reflexão e Crítica, 32(1), 9. 10.1186/s41155-019-0121-8.

Coelho, V. A., Marchante, M., & Romão, A. M. (2022). Adolescents’ trajectories of social anxiety and social withdrawal: Are they influenced by traditional bullying and cyberbullying roles? Contemporary Educational Psychology, 69, 102053. 10.1016/j.cedpsych.2022.102053.

Combs, D. R., Penn, D. L., Wicher, M., & Waldheter, E. (2007). The Ambiguous Intentions Hostility Questionnaire (AIHQ): A new measure for evaluating hostile social-cognitive biases in paranoia. Cognitive Neuropsychiatry, 12(2), 128–143. 10.1080/13546800600787854.

Craske, M. G., Rauch, S. L., Ursano, R., Prenoveau, J., Pine, D. S., & Zinbarg, R. E. (2009). What is an anxiety disorder? Depression and Anxiety, 26(12), 1066–1085. 10.1002/da.20633.

Crick, N. R., & Dodge, K. A. (1994). A review and reformulation of social information-processing mechanisms in children’s social adjustment. Psychological Bulletin, 115(1), 74–101. 10.1037/00332909.115.1.74.

Davidson, R. J., Putnam, K. M., & Larson, C. L. (2000). Dysfunction in the Neural Circuitry of Emotion Regulation–A Possible Prelude to Violence. Science, 289(5479), 591–594. 10.1126/science.289.5479.591.

Demontis, D., Walters, G. B., Athanasiadis, G., Walters, R., Therrien, K., Nielsen, T. T., Farajzadeh, L., Voloudakis, G., Bendl, J., Zeng, B., Zhang, W., Grove, J., Als, T. D., Duan, J., Satterstrom, F. K., Bybjerg-Grauholm, J., Bækved-Hansen, M., Gudmundsson, O. O., Magnusson, S. H., … Børglum, A. D. (2023). Genome-wide analyses of ADHD identify 27 risk loci, refine the genetic architecture and implicate several cognitive domains. Nature Genetics, 55(2), 198–208. 10.1038/s41588-022-01285-8.

Dodge, K. A. (2006). Translational science in action: Hostile attributional style and the development of aggressive behavior problems. Development and Psychopathology, 18(03). 10.1017/S0954579406060391.

Dodge, K. A., Bates, J. E., & Pettit, G. S. (1990). Mechanisms in the Cycle of Violence. Science, 250(4988), 1678–1683. 10.1126/science.2270481.

Dodge, K. A., & Coie, J. D. (1987). Social-Information-Processing Factors in Reactive and Proactive Aggression in Children’s Peer Groups. Journal of Personality and Social Psychology, 13.

Durand, K., Gallay, M., Seigneuric, A., Robichon, F., & Baudouin, J.-Y. (2007). The development of facial emotion recognition: The role of configural information. Journal of Experimental Child Psychology, 97(1), 14–27. 10.1016/j.jecp.2006.12.001.

D’Urso, G., Juvonen, J., & Salmivalli, C. (2023). Do adolescence peer victimization experiences hamper healthy relationships in young adulthood? European Journal of Developmental Psychology, 20(5), 839–853. 10.1080/17405629.2023.2216448.

Forstner, A. J., Awasthi, S., Wolf, C., Maron, E., Erhardt, A., Czamara, D., Eriksson, E., Lavebratt, C., Allgulander, C., Friedrich, N., Becker, J., Hecker, J., Rambau, S., Conrad, R., Geiser, F., McMahon, F. J., Moebus, S., Hess, T., Buerfent, B. C., … Schumacher, J. (2021). Genome-wide association study of panic disorder reveals genetic overlap with neuroticism and depression. Molecular Psychiatry, 26(8), 4179–4190. 10.1038/s41380-019-0590-2.

Fu, X., & Pérez-Edgar, K. (2019). Threat-related attention bias in socioemotional development: A critical review and methodological considerations. Developmental Review, 51, 31–57. 10.1016/j.dr.2018.11.002.

Gallup, A. C., O’Brien, D. T., White, D. D., & Wilson, D. S. (2009). Peer victimization in adolescence has different effects on the sexual behavior of male and female college students. Personality and Individual Differences, 46(5-6), 611–615. 10.1016/j.paid.2008.12.018.

Geoffroy, M.-C., Boivin, M., Arseneault, L., Renaud, J., Perret, L. C., Turecki, G., Michel, G., Salla, J., Vitaro, F., Brendgen, M., Tremblay, R. E., & Côté, S. M. (2018). Childhood trajectories of peer victimization and prediction of mental health outcomes in midadolescence: A longitudinal population-based study. Canadian Medical Association Journal, 190(2), E37–E43. 10.1503/cmaj.170219.

Gibb, B. E., Schofield, C. A., & Coles, M. E. (2009). Reported History of Childhood Abuse and Young Adults’ Information-Processing Biases for Facial Displays of Emotion. Child Maltreatment, 14(2), 148–156. 10.1177/1077559508326358.

Gilbert, P. (2001). EVOLUTION AND SOCIAL ANXIETY. Psychiatric Clinics of North America, 24(4), 723–751. 10.1016/S0193-953X(05)70260-4.

Gini, G., & Pozzoli, T. (2013). Bullied Children and Psychosomatic Problems: A Meta-analysis. Pediatrics, 132(4), 720–729. 10.1542/peds.2013-0614.

Glowacki, L., Wilson, M. L., & Wrangham, R. W. (2020). The evolutionary anthropology of war. Journal of Economic Behavior & Organization, 178, 963–982. 10.1016/j.jebo.2017.09.014.

Grossheinrich, N., Schaeffer, J., Firk, C., Eggermann, T., Huestegge, L., & Konrad, K. (2022). Childhood adversity and approach/avoidance-related behaviour in boys. Journal of Neural Transmission, 129(4), 421–429. 10.1007/s00702-022-02481-w.

Günther, V., Kropidlowski, A., Schmidt, F. M., Koelkebeck, K., Kersting, A., & Suslow, T. (2021). Attentional processes during emotional face perception in social anxiety disorder: A systematic review and meta-analysis of eye-tracking findings. Progress in Neuro-Psychopharmacology and Biological Psychiatry, 111, 110353. 10.1016/j.pnpbp.2021.110353.

Heberlein, A. S., Padon, A. A., Gillihan, S. J., Farah, M. J., & Fellows, L. K. (2008). Ventromedial Frontal Lobe Plays a Critical Role in Facial Emotion Recognition. Journal of Cognitive Neuroscience, 20(4), 721–733. 10.1162/jocn.2008.20049.

Heilbron, N., & Prinstein, M. J. (2008). A Review and Reconceptualization of Social Aggression: Adaptive and Maladaptive Correlates. Clinical Child and Family Psychology Review, 11(4), 176–217. 10.1007/s10567-008-0037-9.

Herd, T., & Kim-Spoon, J. (2021). A Systematic Review of Associations Between Adverse Peer Experiences and Emotion Regulation in Adolescence. Clinical Child and Family Psychology Review, 24(1), 141–163. 10.1007/s10567-020-00337-x.

Hernan, M. A. (2006). Estimating causal effects from epidemiological data. Journal of Epidemiology & Community Health, 60(7), 578–586. 10.1136/jech.2004.029496.

Herzog, S., D’Andrea, W., DePierro, J., & Khedari, V. (2018). When stress becomes the new normal: Alterations in attention and autonomic reactivity in repeated traumatization. Journal of Trauma & Dissociation, 19(3), 362–381. 10.1080/15299732.2018.1441356.

Hiser, J., & Koenigs, M. (2018). The Multifaceted Role of the Ventromedial Prefrontal Cortex in Emotion, Decision Making, Social Cognition, and Psychopathology. Biological Psychiatry, 83(8), 638– 647. 10.1016/j.biopsych.2017.10.030.

Hoepfel, D., Günther, V., Bujanow, A., Kersting, A., Bodenschatz, C. M., & Suslow, T. (2022). Experiences of maltreatment in childhood and attention to facial emotions in healthy young women. Scientific Reports, 12(1), 4317. 10.1038/s41598-022-08290-1.

Ichikawa, H., Nakato, E., Kanazawa, S., Shimamura, K., Sakuta, Y., Sakuta, R., Yamaguchi, M. K., & Kakigi, R. (2014). Hemodynamic response of children with attention-deficit and hyperactive disorder (ADHD) to emotional facial expressions. Neuropsychologia, 63, 51– 58. 10.1016/j.neuropsychologia.2014.08.010.

Iffland, B., Weitkämper, A., Weitkämper, N. J., & Neuner, F. (2019). Attentional avoidance in peer victimized individuals with and without psychiatric disorders. BMC Psychology, 7(1), 12. 10.1186/s40359-019-0284-1.

Ito, T., Yokokawa, K., Yahata, N., Isato, A., Suhara, T., & Yamada, M. (2017). Neural basis of negativity bias in the perception of ambiguous facial expression. Scientific Reports, 7(1), 420. 10.1038/s41598-017-00502-3.

Jaffee, S. R. (2017). Child Maltreatment and Risk for Psychopathology in Childhood and Adulthood. Annual Review of Clinical Psychology, 13(1), 525–551. 10.1146/annurev-clinpsy-032816045005.

Jagannath, V., Grü nblatt, E., Theodoridou, A., Oneda, B., Roth, A., Gerstenberg, M., Franscini, M., Traber-Walker, N., Correll, C. U., Heekeren, K., Rö ssler, W., Rauch, A., & Walitza, S. (2020). Rare copy number variants in individuals at clinical high risk for psychosis: Enrichment of synaptic/brain-related functional pathways. American Journal of Medical Genetics Part B: Neuropsychiatric Genetics, 183(2), 140–151. 10.1002/ajmg.b.32770.

Ke, T., De Simoni, S., Barker, E., & Smith, P. (2022). The association between peer-victimisation and structural and functional brain outcomes: A systematic review. JCPP Advances, 2(2), e12081. 10.1002/jcv2.12081.

Kellij, S., Lodder, G. M. A., Van Den Bedem, N., Güroğlu, B., & Veenstra, R. (2022). The Social Cognitions of Victims of Bullying: A Systematic Review. Adolescent Research Review, 7(3), 287–334. 10.1007/s40894-022-00183-8.

Kindt, M., & Van Den Hout, M. (2001). Selective Attention and Anxiety: A Perspective on Developmental Issues and the Causal Status. Journal of Psychopathology and Behavioral Assessment, 23(3), 193–202. 10.1023/A:1010921405496.

Kircanski, K., Joormann, J., & Gotlib, I. H. (2015). Attention to Emotional Information in Social Anxiety Disorder With and Without Co-Occurring Depression. Cognitive Therapy and Research, 39(2), 153–161. 10.1007/s10608-014-9643-7.

Klein Tuente, S., Bogaerts, S., & Veling, W. (2019). Hostile attribution bias and aggression in adults - a systematic review. Aggression and Violent Behavior, 46, 66–81. 10.1016/j.avb.2019.01.009.

Kochel, K. P., Ladd, G. W., & Rudolph, K. D. (2012). Longitudinal Associations Among Youth Depressive Symptoms, Peer Victimization, and Low Peer Acceptance: An Interpersonal Process Perspective. Child Development, 83(2), 637–650. 10.1111/j.1467-8624.2011.01722.x.

Kruglanski, A. W., Ellenberg, M., Szumowska, E., Molinario, E., Speckhard, A., Leander, N. P., Pierro, A., Di Cicco, G., & Bushman, B. J. (2023). Frustration–aggression hypothesis reconsidered: The role of significance quest. Aggressive Behavior, 49(5), 445–468. 10.1002/ab.22092.

Lisk, S., Vaswani, A., Linetzky, M., Bar-Haim, Y., & Lau, J. Y. (2020). Systematic Review and Meta-Analysis: Eye-Tracking of Attention to Threat in Child and Adolescent Anxiety. Journal of the American Academy of Child & Adolescent Psychiatry, 59(1), 88–99.e1. 10.1016/j.jaac.2019.06.006.

Liu, Y., Wang, Y., Gozli, D. G., Xiang, Y.-T., & Jackson, T. (2021). Current status of the anger superiority hypothesis: A meta-analytic review of N2pc studies. Psychophysiology, 58(1), e13700. 10.1111/psyp.13700.

Ma, D. S., Correll, J., & Wittenbrink, B. (2015). The Chicago face database: A free stimulus set of faces and norming data. Behavior Research Methods, 47(4), 1122–1135. 10.3758/s13428-014-0532-5.

MacLeod, C., Mathews, A., & Tata, P. (1986). Attentional bias in emotional disorders. Journal of Abnormal Psychology, 95(1), 15–20. 10.1037/0021-843X.95.1.15.

Maoz, K., Eldar, S., Stoddard, J., Pine, D. S., Leibenluft, E., & Bar-Haim, Y. (2016). Angry-happy interpretations of ambiguous faces in social anxiety disorder. Psychiatry Research, 241, 122–127. 10.1016/j.psychres.2016.04.100.

McLaughlin, K. A., Peverill, M., Gold, A. L., Alves, S., & Sheridan, M. A. (2015). Child Maltreatment and Neural Systems Underlying Emotion Regulation. Journal of the American Academy of Child & Adolescent Psychiatry, 54(9), 753–762. 10.1016/j.jaac.2015.06.010.

Monk, C. S., Nelson, E. E., McClure, E. B., Mogg, K., Bradley, B. P., Leibenluft, E., Blair, R. J. R., Chen, G., Charney, D. S., Ernst, M., & Pine, D. S. (2006). Ventrolateral Prefrontal Cortex Activation and Attentional Bias in Response to Angry Faces in Adolescents With Generalized Anxiety Disorder. American Journal of Psychiatry, 163(6), 1091–1097. 10.1176/ajp.2006.163.6.1091.

Murray, A. L., Eisner, M., Ribeaud, D., Kaiser, D., McKenzie, K., & Murray, G. (2021). Validation of a Brief Self-Report Measure of Adolescent Bullying Perpetration and Victimization [Publisher: SAGE Publications]. Assessment, 28(1), 128–140. 10.1177/1073191119858406.

Norman, R. E., Byambaa, M., De, R., Butchart, A., Scott, J., & Vos, T. (2012). The Long-Term Health Consequences of Child Physical Abuse, Emotional Abuse, and Neglect: A Systematic Review and Meta-Analysis (M. Tomlinson, Ed.). PLoS Medicine, 9(11), e1001349. 10.1371/journal.pmed.1001349.

O’Mahen, H. A., Karl, A., Moberly, N., & Fedock, G. (2015). The association between childhood maltreatment and emotion regulation: Two different mechanisms contributing to depression? Journal of Affective Disorders, 174, 287–295. 10.1016/j.jad.2014.11.028.

Ossa, F. C., Pietrowsky, R., Bering, R., & Kaess, M. (2019). Symptoms of posttraumatic stress disorder among targets of school bullying. Child and Adolescent Psychiatry and Mental Health, 13(1), 43. 10.1186/s13034-019-0304-1.

Otowa, T., Hek, K., Lee, M., Byrne, E. M., Mirza, S. S., Nivard, M. G., Bigdeli, T., Aggen, S. H., Adkins, D., Wolen, A., Fanous, A., Keller, M. C., Castelao, E., Kutalik, Z., Der Auwera, S. V., Homuth, G., Nauck, M., Teumer, A., Milaneschi, Y., … Hettema, J. M. (2016). Meta-analysis of genome-wide association studies of anxiety disorders. Molecular Psychiatry, 21(10), 1391–1399. 10.1038/mp.2015.197.

Pantoja-Urbán, A. H., Richer, S., Mittermaier, A., Giroux, M., Nouel, D., Hernandez, G., & Flores, C. (2024). Gains and Losses: Resilience to Social Defeat Stress in Adolescent Female Mice. Biological Psychiatry, 95(1), 37–47. 10.1016/j.biopsych.2023.06.014.

Penton-Voak, I. S., Thomas, J., Gage, S. H., McMurran, M., McDonald, S., & Munafò, M.R. (2013). Increasing Recognition of Happiness in Ambiguous Facial Expressions Reduces Anger and Aggressive Behavior. Psychological Science, 24(5), 688–697. 10.1177/0956797612459657.

Perren, S., Ettekal, I., & Ladd, G. (2013). The impact of peer victimization on later maladjustment: Mediating and moderating effects of hostile and self-blaming attributions. Journal of Child Psychology and Psychiatry, 54(1), 46–55. 10.1111/j.14697610.2012.02618.x.

Pfaltz, M. C., Passardi, S., Auschra, B., Fares-Otero, N. E., Schnyder, U., & Peyk, P. (2019). Are you angry at me? Negative interpretations of neutral facial expressions are linked to child maltreatment but not to posttraumatic stress disorder. European Journal of Psychotraumatology, 10(1), 1682929. 10.1080/20008198.2019.1682929.

Pfattheicher, S., & Keller, J. (2014). Towards a Biopsychological Understanding of Costly Punishment: The Role of Basal Cortisol (A. Szolnoki, Ed.). PLoS ONE, 9(1), e85691. 10.1371/journal.pone.0085691.

Pine, D. S., Mogg, K., Bradley, B. P., Montgomery, L., Monk, C. S., McClure, E., Guyer, A. E., Ernst, M., Charney, D. S., & Kaufman, J. (2005). Attention Bias to Threat in Maltreated Children: Implications for Vulnerability to Stress-Related Psychopathology. American Journal of Psychiatry, 162(2), 291–296. 10.1176/appi.ajp.162.2.291.

Plana, I., Lavoie, M.-A., Battaglia, M., & Achim, A. M. (2014). A meta-analysis and scoping review of social cognition performance in social phobia, posttraumatic stress disorder and other anxiety disorders. Journal of Anxiety Disorders, 28(2), 169–177. 10.1016/j.janxdis.2013.09.005.

Pollak, S. D., & Kistler, D. J. (2002). Early experience is associated with the development of categorical representations for facial expressions of emotion. Proceedings of the National Academy of Sciences, 99(13), 9072–9076. 10.1073/pnas.142165999.

Pontillo, M., Tata, M. C., Averna, R., Demaria, F., Gargiullo, P., Guerrera, S., Pucciarini, M. L., Santonastaso, O., & Vicari, S. (2019). Peer Victimization and Onset of Social Anxiety Disorder in Children and Adolescents. Brain Sciences, 9(6), 132. 10.3390/brainsci9060132.

Price, A. L., Patterson, N. J., Plenge, R. M., Weinblatt, M. E., Shadick, N. A., & Reich, D. (2006). Principal components analysis corrects for stratification in genome-wide association studies. Nature Genetics, 38(8), 904–909. 10.1038/ng1847.

Purcell, S., Neale, B., Todd-Brown, K., Thomas, L., Ferreira, M. A., Bender, D., Maller, J., Sklar, P., De Bakker, P. I., Daly, M. J., & Sham, P. C. (2007). PLINK: A Tool Set for Whole-Genome Association and Population-Based Linkage Analyses. The American Journal of Human Genetics, 81(3), 559–575. 10.1086/519795.

Rapee, R. M., Creswell, C., Kendall, P. C., Pine, D. S., & Waters, A. M. (2023). Anxiety disorders in children and adolescents: A summary and overview of the literature. Behaviour Research and Therapy, 168, 104376. 10.1016/j.brat.2023.104376.

Ribeaud, D., Murray, A., Shanahan, L., Shanahan, M. J., & Eisner, M. (2022). Cohort Profile: The Zurich Project on the Social Development from Childhood to Adulthood (z-proso). Journal of Developmental and Life-Course Criminology, 8(1), 151–171. 10.1007/s40865-022-00195-x.

Riley, W. T., Treiber, F. A., & Woods, M. G. (1989). Anger and Hostility in Depression: The Journal of Nervous and Mental Disease, 177(11), 668–674. 10.1097/00005053-198911000-00002.

Rosenbaum, P. R., & Rubin, D. B. (1984). Reducing Bias in Observational Studies Using Subclassification on the Propensity Score. Journal of the American Statistical Association, 79(387), 516–524. 10.2307/2288398.

Roy, A. K., Vasa, R. A., Bruck, M., Mogg, K., Bradley, B. P., Sweeney, M., Bergman, R. L., Mcclure-Tone, E. B., & Pine, D. S. (2008). Attention Bias Toward Threat in Pediatric Anxiety Disorders. Journal of the American Academy of Child & Adolescent Psychiatry, 47(10), 1189–1196. 10.1097/CHI.0b013e3181825ace.

Rubin, D. B. (1976). Inference and missing data. Biometrika, 63(3), 581–592. 10.1093/biomet/63.3.581.

Rubin, D. B. (1997). Estimating Causal Effects from Large Data Sets Using Propensity Scores. Annals of Internal Medicine, 8(2), 757–63. 10.7326/0003-4819-127-8_part_2-199710151-00064.

Rubin, K. H., Coplan, R. J., & Bowker, J. C. (2009). Social Withdrawal in Childhood. Annual Review of Psychology, 60(1), 141–171. 10.1146/annurev.psych.60.110707.163642.

Rutter, M., Pickles, A., Murray, R., & Eaves, L. (2001). Testing hypotheses on specific environmental causal effects on behavior. Psychological Bulletin, 127(3), 291–324. 10.1037//0033-2909.127.3.291.

Samson, J. A., Newkirk, T. R., & Teicher, M. H. (2024). Practitioner Review: Neurobiological consequences of childhood maltreatment – clinical and therapeutic implications for practitioners. Journal of Child Psychology and Psychiatry, 65(3), 369–380. 10.1111/jcpp.13883.

Sapolsky, R. M. (2004). Social Status and Health in Humans and Other Animals. Annual Review of Anthropology, 33(1), 393–418. 10.1146/annurev.anthro.33.070203.144000.

Sato, W., Kubota, Y., Okada, T., Murai, T., Yoshikawa, S., & Sengoku, A. (2002). Seeing Happy Emotion in Fearful and Angry Faces: Qualitative Analysis of Facial Expression Recognition in a Bilateral Amygdala-Damaged Patient. Cortex, 38(5), 727–742. 10.1016/S0010-9452(08)70040-6.

Schacter, H. L., Marusak, H. A., Borg, B. A., & Jovanovic, T. (2024). Facing ambiguity: Social threat sensitivity mediates the association between peer victimization and adolescent anxiety. Development and Psychopathology, 36(1), 112–120. 10.1017/S0954579422001018.

Schimmack, U. (2005). Attentional Interference Effects of Emotional Pictures: Threat, Negativity, or Arousal? (D. Derryberry, Ed.). Emotion, 5(1), 55–66. 10.1037/1528-3542.5.1.55.

Shechner, T., Britton, J. C., Pérez-Edgar, K., Bar-Haim, Y., Ernst, M., Fox, N. A., Leibenluft, E., & Pine, D. S. (2012). Attention biases, anxiety, and development: Toward or away from threats or rewards? Depression and Anxiety, 29(4), 282–294. 10.1002/da.20914.

Siegel, R. S., La Greca, A. M., & Harrison, H. M. (2009). Peer Victimization and Social Anxiety in Adolescents: Prospective and Reciprocal Relationships. Journal of Youth and Adolescence, 38(8), 1096–1109. 10.1007/s10964-009-9392-1.

Silberg, J. L., Copeland, W., Linker, J., Moore, A. A., Roberson-Nay, R., & York, T. P. (2016). Psychiatric outcomes of bullying victimization: A study of discordant monozygotic twins. Psychological Medicine, 46(9), 1875–1883. 10.1017/S0033291716000362.

Slavich, G. M. (2020). Social Safety Theory: A Biologically Based Evolutionary Perspective on Life Stress, Health, and Behavior. Annual Review of Clinical Psychology, 16(1), 265–295. 10.1146/annurev-clinpsy-032816-045159.

Stapinski, L. A., Bowes, L., Wolke, D., Pearson, R. M., Mahedy, L., Button, K. S., Lewis, G., & Araya, R. (2014). Peer Victimization During Adolescence and Risk For Anxiety Disorders in Adulthood: A Prospective Cohort Study. Depression and Anxiety, 31(7), 574–582. 10.1002/da.22270.

Stekhoven, D. J., & Buhlmann, P. (2012). MissForest—non-parametric missing value imputation for mixed-type data. Bioinformatics, 28(1), 112–118. 10.1093/bioinformatics/btr597.

Stilling, R. M., Moloney, G. M., Ryan, F. J., Hoban, A. E., Bastiaanssen, T. F., Shanahan, F., Clarke, G., Claesson, M. J., Dinan, T. G., & Cryan, J. F. (2018). Social interaction-induced activation of RNA splicing in the amygdala of microbiome-deficient mice. eLife, 7. 10.7554/eLife.33070.

Taylor, S. E., Eisenberger, N. I., Saxbe, D., Lehman, B. J., & Lieberman, M. D. (2006). Neural Responses to Emotional Stimuli Are Associated with Childhood Family Stress. Biological Psychiatry, 60(3), 296–301. 10.1016/j.biopsych.2005.09.027.

Tremblay, R. E. (1994). Predicting Early Onset of Male Antisocial Behavior From Preschool Behavior. Archives of General Psychiatry, 51(9), 732. 10.1001/archpsyc.1994.03950090064009.

Tsuchida, A., & Fellows, L. K. (2012). Are You Upset? Distinct Roles for Orbitofrontal and Lateral Prefrontal Cortex in Detecting and Distinguishing Facial Expressions of Emotion. Cerebral Cortex, 22(12), 2904–2912. 10.1093/cercor/bhr370.

Van Bockstaele, B., Verschuere, B., Tibboel, H., De Houwer, J., Crombez, G., & Koster, E. H. W. (2014). A review of current evidence for the causal impact of attentional bias on fear and anxiety. Psychological Bulletin, 140(3), 682–721. 10.1037/a0034834.

Ver Hoeve, E., Kelly, G., Luz, S., Ghanshani, S., & Bhatnagar, S. (2013). Short-term and long-term effects of repeated social defeat during adolescence or adulthood in female rats. Neuroscience, 249, 63–73. 10.1016/j.neuroscience.2013.01.073.

Verhoef, R. E., Alsem, S. C., Verhulp, E. E., & De Castro, B. O. (2019). Hostile Intent Attribution and Aggressive Behavior in Children Revisited: A Meta-Analysis. Child Development, 90(5). 10.1111/cdev.13255.

Volk, A. A., Dane, A. V., & Al-Jbouri, E. (2022). Is Adolescent Bullying an Evolutionary Adaptation? A 10-Year Review. Educational Psychology Review, 34(4), 2351–2378. 10.1007/s10648022-09703-3.

Wald, I., Lubin, G., Holoshitz, Y., Muller, D., Fruchter, E., Pine, D. S., Charney, D. S., & Bar-Haim, Y. (2011). Battlefield-like stress following simulated combat and suppression of attention bias to threat. Psychological Medicine, 41(4), 699–707. 10.1017/S0033291710002308.

Wilkowski, B. M., & Robinson, M. D. (2008). The Cognitive Basis of Trait Anger and Reactive Aggression: An Integrative Analysis. Personality and Social Psychology Review, 12(1), 3–21. 10.1177/1088868307309874.

Williams, K. C., Nto, N. J., Van Vuren, E. J., Sallie, F. N., Molebatsi, K., Kroneberg, K. S., Roomaney, A. A., Salie, M., & Womersley, J. S. (2024). Early biological and psychosocial factors associated with PTSD onset and persistence in youth. European Journal of Psychotraumatology, 15(1), 2432160. 10.1080/20008066.2024.2432160.

Wormwood, J. B., Lynn, S. K., Feldman Barrett, L., & Quigley, K. S. (2016). Threat perception after the Boston Marathon bombings: The effects of personal relevance and conceptual framing. Cognition and Emotion, 30(3), 539–549. 10.1080/02699931.2015.1010487.

Wray, N. R., Ripke, S., Mattheisen, M., Trzaskowski, M., Byrne, E. M., Abdellaoui, A., Adams, M. J., Agerbo, E., Air, T. M., Andlauer, T. M. F., Bacanu, S.-A., Bækvad-Hansen, M., Beekman, A. F. T., Bigdeli, T. B., Binder, E. B., Blackwood, D. R. H., Bryois, J., Buttenschøn, H. N., Bybjerg-Grauholm, J., … Sullivan, P. F. (2018). Genome-wide association analyses identify 44 risk variants and refine the genetic architecture of major depression. Nature Genetics, 50(5), 668–681. 10.1038/s41588-018-0090-3.

Ziv, Y., Leibovich, I., & Shechtman, Z. (2013). Bullying and victimization in early adolescence: Relations to social information processing patterns. Aggressive Behavior, 39(6), 482–492. 10.1002/ab.21494.

z-proso Project Team. (2024). Z-proso Handbook: Instruments and Procedures in the Adolescent and Young Adult Surveys (Age 11 to 24; Waves K4-K9) [Publisher: Jacobs Center for Productive Youth Development, University of Zurich]. 10.5167/UZH253680.

Zuber, S., Bechtiger, L., Bodelet, J. S., Golin, M., Heumann, J., Kim, J. H., Klee, M., Mur, J., Noll, J., Voll, S., O’Keefe, P., Steinhoff, A., Zö litz, U., Muniz-Terrera, G., Shanahan, L., Shanahan, M. J., & Hofer, S. M. (2023). An integrative approach for the analysis of risk and health across the life course: Challenges, innovations, and opportunities for life course research. Discover Social Science and Health, 3(1), 14. 10.1007/s44155-023-00044-2.

Zupan, B., & Eskritt, M. (2024). Facial and Vocal Emotion Recognition in Adolescence: A Systematic Review. Adolescent Research Review, 9(2), 253–277. 10.1007/s40894-023-00219-7.

## Supporting references

Austin, P. C., & Stuart, E. A. (2017). The performance of inverse probability of treatment weighting and full matching on the propensity score in the presence of model misspecification when estimating the effect of treatment on survival outcomes. Statistical Methods in Medical Research, 26(4), 1654–1670. 10.1177/0962280215584401.

Brendgen, M., Girard, A., Vitaro, F., Dionne, G., & Boivin, M. (2015). Gene-Environment Correlation Linking Aggression and Peer Victimization: Do Classroom Behavioral Norms Matter? Journal of Abnormal Child Psychology, 43(1), 19–31. 10.1007/s10802-013-9807-z.

Casper, D. M., & Card, N. A. (2017). Overt and Relational Victimization: A Meta-Analytic Review of Their Overlap and Associations With Social-Psychological Adjustment. Child Development, 88(2), 466–483. 10.1111/cdev.12621.

Cattell, R. B. (1966). The Scree Test For The Number Of Factors. Multivariate Behavioral Research, 1(2), 245–276. 10.1207/s15327906mbr0102_10.

Cook, C. R., Williams, K. R., Guerra, N. G., Kim, T. E., & Sadek, S. (2010). Predictors of bullying and victimization in childhood and adolescence: A meta-analytic investigation. School Psychology Quarterly, 25(2), 65–83. 10.1037/a0020149.

Dantchev, S., & Wolke, D. (2019). Trouble in the nest: Antecedents of sibling bullying victimization and perpetration. Developmental Psychology, 55(5), 1059–1071. 10.1037/dev0000700.

Davison, A. C., & Hinkley, D. V. (1997). Bootstrap Methods and their Application. Cambridge University Press.

Dede, B., Delk, L., & White, B. A. (2021). Relationships between facial emotion recognition, internalizing symptoms, and social problems in young children. Personality and Individual Differences, 171, 110448. 10.1016/j.paid.2020.110448.

Delprato, M., Akyeampong, K., & Dunne, M. (2017). The impact of bullying on students’ learning in Latin America: A matching approach for 15 countries. International Journal of Educational Development, 52, 37–57. 10.1016/j.ijedudev.2016.10.002.

DiLalla, L. F., Jamnik, M. R., Marshall, R. L., Weisbecker, R., & Vazquez, C. (2021). Birth Complications and Negative Emotionality Predict Externalizing Behaviors in Young Twins: Moderations with Genetic and Family Risk Factors. Behavior Genetics, 51(5), 463–475. 10.1007/s10519-021-10062-y.

Duffy, A. L., & Nesdale, D. (2009). Peer Groups, Social Identity, and Children’s Bullying Behavior. Social Development, 18(1), 121–139. 10.1111/j.1467-9507.2008.00484.x.

Eastman, M. L., Verhulst, B., Rappaport, L. M., Dirks, M., Sawyers, C., Pine, D. S., Leibenluft, E., Brotman, M. A., Hettema, J. M., & Roberson-Nay, R. (2018). Age-Related Differences in the Structure of Genetic and Environmental Contributions to Types of Peer Victimization. Behavior Genetics, 48(6), 421–431. 10.1007/s10519-018-9923-1.

Espelage, D. L., Bosworth, K., & Simon, T. R. (2001). Short-Term Stability and Prospective Correlates of Bullying in Middle-School Students: An Examination of Potential Demographic, Psychosocial, and Environmental Influences. Violence and Victims, 16(4), 411–426. 10.1891/0886-6708.16.4.411.

Forster, M., Gower, A. L., McMorris, B. J., & Borowsky, I. W. (2020). Adverse Childhood Experiences and School-Based Victimization and Perpetration. Journal of Interpersonal Violence, 35(3-4), 662–681. 10.1177/0886260517689885.

Hernán, M. A., & Robins, J. M. (2020). Causal Inference: What If, 311.

Holt, M. K., & Espelage, D. L. (2007). Perceived Social Support among Bullies, Victims, and Bully-Victims. Journal of Youth and Adolescence, 36(8), 984–994. 10.1007/s10964-006-9153-3.

Jennings, W. G., Park, M., Tomsich, E. A., Gover, A. R., & Akers, R. L. (2011). Assessing the Overlap in Dating Violence Perpetration and Victimization among South Korean College Students: The Influence of Social Learning and SelfControl. American Journal of Criminal Justice, 36(2), 188– 206. 10.1007/s12103-011-9110-x.

Karišik, M. (2019). Postoperative Pain and Stress Response: Does Child’s Gender Have an Influence? Acta Clinica Croatica. 10.20471/acc.2019.58.02.10.

Kemeny, M. E. (2003). The Psychobiology of Stress. Current Directions in Psychological Science, 12(4), 124–129. 10.1111/1467-8721.01246.

Lereya, S. T., Copeland, W. E., Costello, E. J., & Wolke, D. (2015). Adult mental health consequences of peer bullying and maltreatment in childhood: Two cohorts in two countries. The Lancet Psychiatry, 2(6), 524–531. 10.1016/S2215-0366(15)00165-0.

Lereya, S. T., Samara, M., & Wolke, D. (2013). Parenting behavior and the risk of becoming a victim and a bully/victim: A meta-analysis study. Child Abuse & Neglect, 37(12), 1091–1108. 10.1016/j.chiabu.2013.03.001.

Martínez, I., Murgui, S., Garcia, O. F., & Garcia, F. (2019). Parenting in the digital era: Protective and risk parenting styles for traditional bullying and cyberbullying victimization. Computers in Human Behavior, 90, 84–92. 10.1016/j.chb.2018.08.036.

Morgan, S. L., & Winship, C. (2015). Counterfactuals and causal inference. methods and principles for social research. (Second Edition). Cambridge University Press.

Murray, A. L., Obsuth, I., Eisner, M., & Ribeaud, D. (2019). Evaluating Longitudinal Invariance in Dimensions of Mental Health Across Adolescence: An Analysis of the Social Behavior Questionnaire [Publisher: SAGE Publications]. Assessment, 26(7), 1234–1245. 10.1177/1073191117721741.

Penner-Goeke, S., Bothe, M., Rek, N., Kreitmaier, P., Pö hlchen, D., Kühnel, A., Glaser, L. V., Kaya, E., Krontira, A. C., Röh, S., Czamara, D., Kö del, M., Monteserin-Garcia, J., Diener, L., Wö lfel, B., Sauer, S., Rummel, C., Riesenberg, S., Arloth-Knauer, J., … Binder, E. B. (2023). High-throughput screening of glucocorticoid-induced enhancer activity reveals mechanisms of stress-related psychiatric disorders. Proceedings of the National Academy of Sciences, 120(49), e2305773120. 10.1073/pnas.2305773120.

Perhamus, G. R., Perry, K. J., Murray-Close, D., & Ostrov, J. M. (2022). Stress reactivity and social cognition in pure and co-occurring early childhood relational bullying and victimization. Development and Psychopathology, 34(4), 1300–1312. 10.1017/S0954579421000298.

Price, A. L., Patterson, N. J., Plenge, R. M., Weinblatt, M. E., Shadick, N. A., & Reich, D. (2006). Principal components analysis corrects for stratification in genomewide association studies. Nature Genetics, 38(8), 904–909. 10.1038/ng1847.

Puhl, R. M., Luedicke, J., & Heuer, C. (2011). Weight-Based Victimization Toward Overweight Adolescents: Observations and Reactions of Peers. Journal of School Health, 81(11), 696–703. 10.1111/j.1746-1561.2011.00646.x.

Qin, D. B., Way, N., & Rana, M. (2008). The “model minority” and their discontent: Examining peer discrimination and harassment of Chinese American immigrant youth. New Directions for Child and Adolescent Development, 2008(121), 27–42. 10.1002/cd.221.

Reifeis, S. A., & Hudgens, M. G. (2022). On Variance of the Treatment Effect in the Treated When Estimated by Inverse Probability Weighting. American Journal of Epidemiology, 191(6), 1092– 1097. 10.1093/aje/kwac014.

Robson, D. A., Allen, M. S., & Howard, S. J. (2020). Self-regulation in childhood as a predictor of future outcomes: A metaanalytic review. Psychological Bulletin, 146(4), 324–354. 10.1037/bul0000227.

Rose, C. A., Simpson, C. G., & Preast, J. L. (2016). Exploring Psychosocial Predictors of Bullying Involvement for Students With Disabilities. Remedial and Special Education, 37(5), 308–317. 10.1177/0741932516629219.

Schoeler, T., Duncan, L., Cecil, C. M., Ploubidis, G. B., & Pingault, J.-B. (2018). Quasi-experimental evidence on short- and long-term consequences of bullying victimization: A meta-analysis. Psychological Bulletin, 144(12), 1229–1246. 10.1037/bul0000171.

Schultz, D., Izard, C. E., & Bear, G. (2004). Children’s emotion processing: Relations to emotionality and aggression. Development and Psychopathology, 16(02). 10.1017/S0954579404044566.

Shanahan, M. J., Cole, S. W., Ravi, S., Chumbley, J., Xu, W., Potente, C., Levitt, B., Bodelet, J., Aiello, A., Gaydosh, L., & Harris, K. M. (2022). Socioeconomic inequalities in molecular risk for chronic diseases observed in young adulthood. Proceedings of the National Academy of Sciences, 119(43), e2103088119. 10.1073/pnas.2103088119.

Stuart, E. A. (2010). Matching Methods for Causal Inference: A Review and a Look Forward. Statistical Science, 25(1), 1–21. 10.1214/09-STS313.

Tippett, N., & Wolke, D. (2014). Socioeconomic Status and Bullying: A Meta-Analysis. American Journal of Public Health, 104(6), e48–e59. 10.2105/AJPH.2014.301960.

White, J. D., & Kaffman, A. (2020). Editorial Perspective: Childhood maltreatment – the problematic unisex assumption. Journal of Child Psychology and Psychiatry, 61(6), 732–734. 10.1111/jcpp.13177.

Wong, J. S., & Schonlau, M. (2013). Does Bully Victimization Predict Future Delinquency?: A Propensity Score Matching Approach. Criminal Justice and Behavior, 40(11), 1184–1208. 10.1177/0093854813503443.

Wu, X., Zhen, R., Shen, L., Tan, R., & Zhou, X. (2023). Patterns of Elementary School Students’ Bullying Victimization: Roles of Family and Individual Factors. Journal of Interpersonal Violence, 38(3-4), 2410–2431. 10.1177/08862605221101190.

z-proso Project Team. (2024). Z-proso Handbook: Instruments and Procedures in the Adolescent and Young Adult Surveys (Age 11 to 24; Waves K4-K9) [Publisher: Jacobs Center for Productive Youth Development, University of Zurich]. 10.5167/UZH-253680.

Zych, I., Farrington, D. P., Llorent, V. J., Ribeaud, D., & Eisner, M. P. (2020). Childhood Risk and Protective Factors as Predictors of Adolescent Bullying Roles. International Journal of Bullying Prevention. 10.1007/s42380-020-00068-1.

